# Epigenetic Instability May Alter Cell State Transitions and Anticancer Drug Resistance

**DOI:** 10.1101/2020.11.15.383521

**Authors:** Anshul Saini, James M. Gallo

**Affiliations:** Department of Pharmaceutical Sciences, University at Buffalo, Buffalo, NY

## Abstract

Drug resistance is a significant obstacle to successful and durable anti-cancer therapy. Targeted therapy is often effective during early phases of treatment; however, eventually cancer cells adapt and transition to drug-resistant cells states rendering the treatment ineffective. It is proposed that cell state can be a determinant of drug efficacy and manipulated to affect the development of anticancer drug resistance. In this work, we developed two stochastic cell state models – referenced to brain tumors - that included transcriptionally-permissive and -restrictive states based on the underlying hypothesis that epigenetic instability mitigates lock-in of drug-resistant states. One model used single-step state transitions, whereas the other considered a multi-step process to lock-in drug resistance. The latter model showed that with moderate epigenetic instability the drug-resistant cell populations were reduced, on average, by 60%, whereas a high level of epigenetic disruption reduced them by about 90%. Generation of epigenetic instability via epigenetic modifier therapy could be a viable strategy to mitigate anticancer drug resistance.

## Introduction

The hallmarks of cancer have evolved from their original inception and indicate the complexity of the disease and the remarkable adaptability of cancer cells to sustain growth under adverse conditions including drug therapy [1, 2]. Epigenetic reprogramming or plasticity is part of the adaptability armamentarium of cancer cells that alters gene expression by regulating gene transcription through histone post-translational modifications (PTMs). Thus, disrupted epigenetic mechanisms play a pivotal role in cancer biology that are characterized by epigenetic plasticity and altered cell states [3, 4]. In recent years, epigenetic modifiers have been used in attempts to reverse resistance to other drugs through epigenetic modifications [5–10]. The reprogramming capability of the epigenome indicates that cells are susceptible to disequilibrium and it is precisely this instability that may be tapped to offer a therapeutic strategy to improve drug therapy.

We previously presented a deterministic cell state model of mutant isocitrate dehydrogenase-1 (IDH1) gliomas that consisted of quiescent, stem and differentiated glioma cells that could transition between each other reflecting intratumoral heterogeneity (ITH) and cellular adaptation [11]. The model incorporated cell proliferation, death, and state transitions in the context of drug resistance. Model simulations indicated that modulation of cell transition rates based upon fluctuating D-2-hydroxy-glutatarate (D2HG) concentrations, a known epigenetic modifier of this type of brain tumor, partially mitigated the evolution of drug-resistant tumors. However, the intrinsic randomness of cell biology is more appropriately captured by stochastic modeling approaches. Various studies have shown that models based on Markovian stochastic transitions between tumor cell states are consistent with observed attributes of tumor heterogeneity [12–14]. In this investigation, the stochastic cell state models consider epigenetic-mediated transcriptionally-permissive and transcriptionally-restrictive states as precursors to drug resistant states. The mathematical models show that implementation of epigenetic instability may be a strategy to subdue the evolution of drug resistance.

## Results

### Basic Single-Step Model

A solid tumor, and herein referred to as a brain tumor or glioblastoma multiforme (GBM), was considered to consist of multiple cell types or states (Figure 1). The cell populations change over time due to proliferation, death, and state transitions. State transitions may be initiated due to the microenvironment (blood flow and hypoxia) and drug therapy as a means to survive and grow. In the model (Figure 1), at time zero, there are five cell states wherein both differentiated glioma (G) and glioma stem cells (GS) exist in two transcriptional states, either permissive (Tp) or restrictive (Tr) that leads to four cell states (G-Tp, G-Tr, GS-Tp, and GS-Tr).

**Figure 1:**
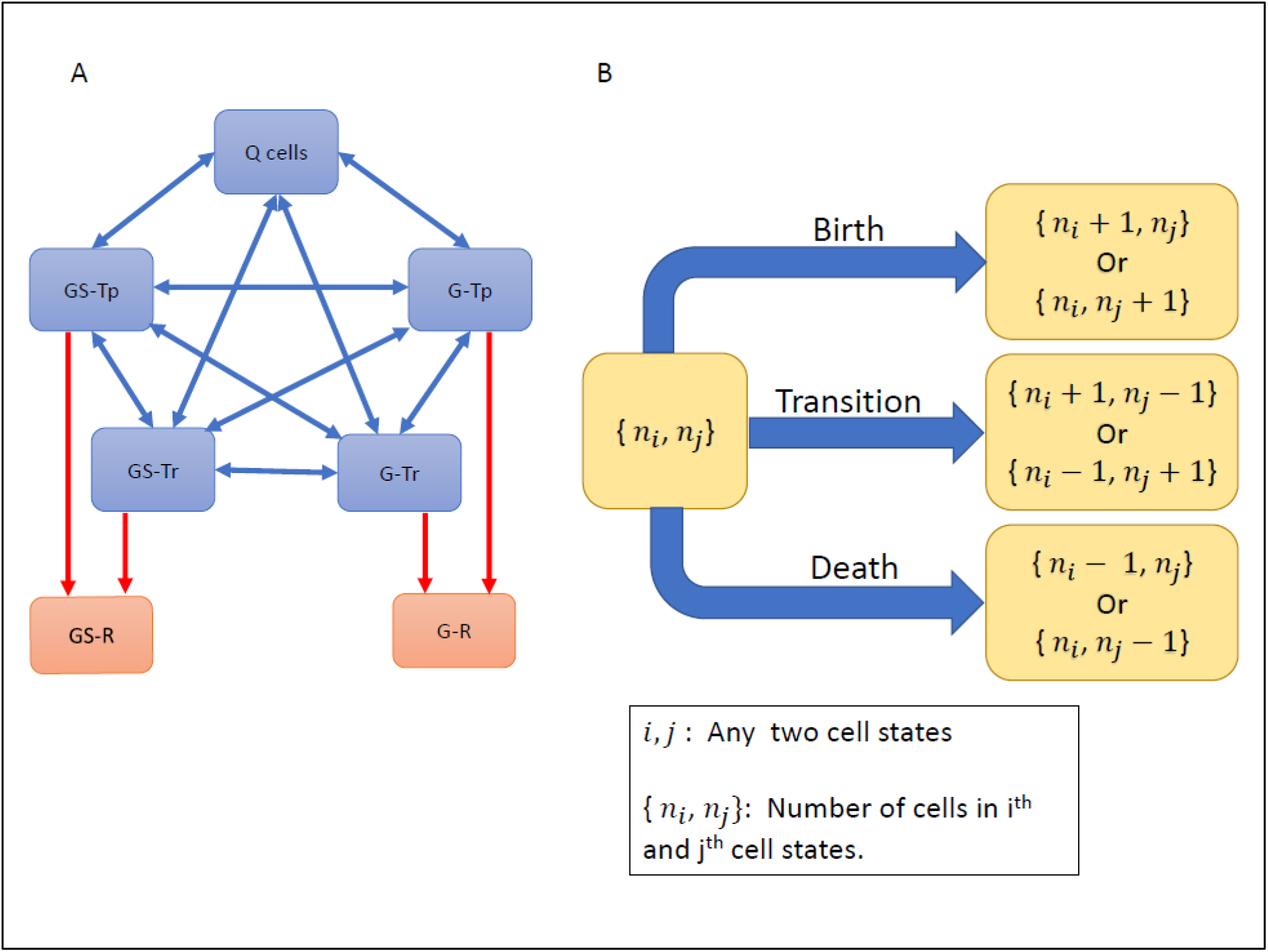
Schematic of the basic single-step cell state model. (A) The model consists of five states: Quiescent cells (Q), transcriptionally-permissive Glioma and Glioma Stem cells (G-Tp and GS-Tp respectively), and transcriptionally-restrictive glioma and glioma stem cells (G-Tr and GS-Tr respectively). Cells can transition between any two of these states (indicated by two-sided blue arrows). In addition, in the presence of drug therapy, cells may transition to drug resistant states (G-R/GS-R). Once in a resistant cell state, the cells are unable to transition back to drug-sensitive states (indicated by one-way red arrows). (B) Cells can undergo birth (except Q cells), death or transition to another state.

Quiescent (Q) cells comprise a fifth cell state. Drug therapy can also lead to two more cell states, drug resistant glioma (G-R) and glioma stem cells (GS-R). Cells in a particular state can undergo either birth, death or transition to another cell state, except Q cells that may only die or transition. This construction of the model permits an understanding of how epigenetic changes affect cell state dynamics. The first set of simulations are shown in Figure 2 using the parameters listed in Table 1 that indicate birth, death and cell transition rates for each cell type under different conditions.

**Figure 2:**
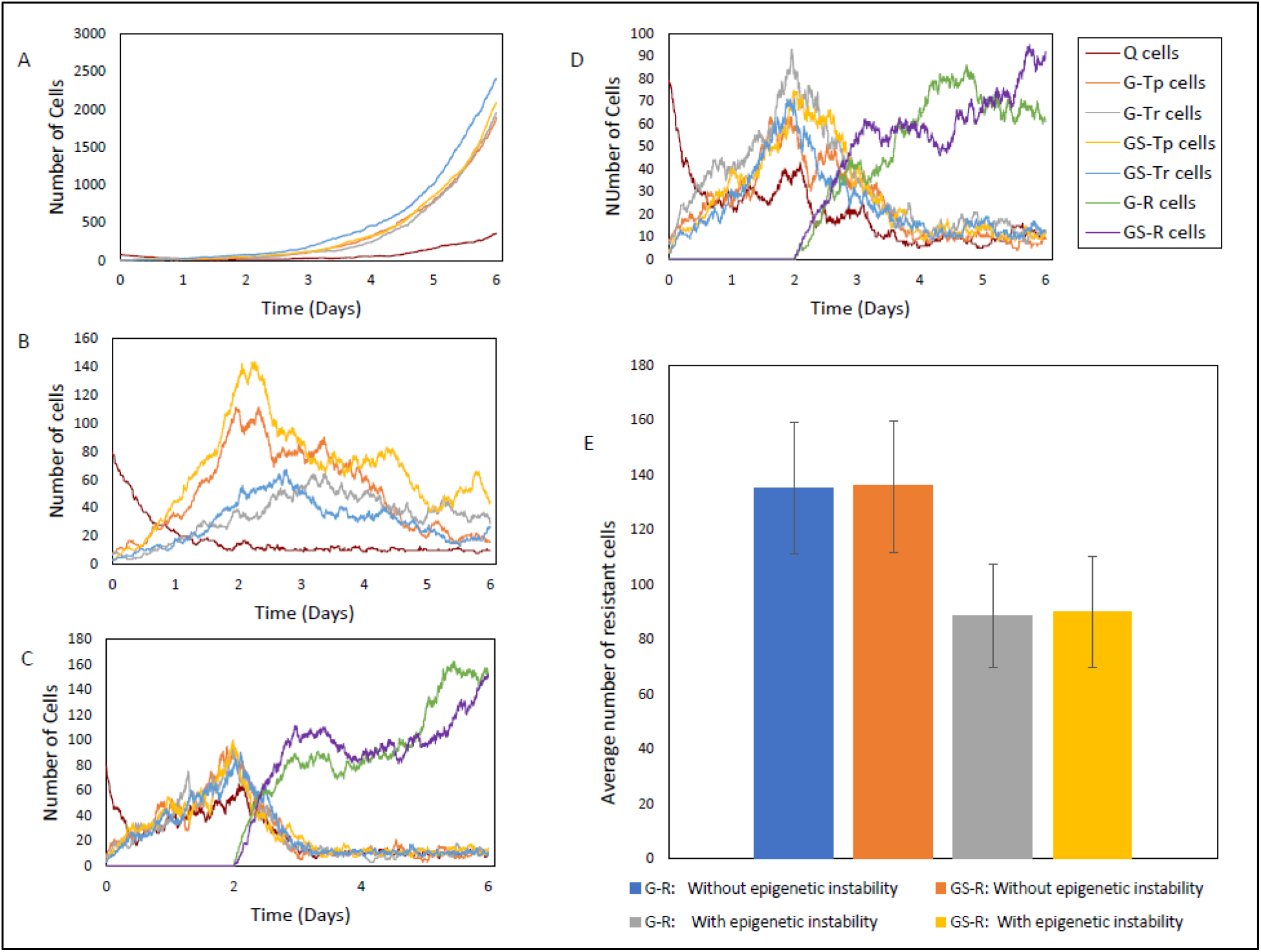
Cell population dynamics for the basic single-step model. (A) Control with low transition rates and no drug exposure. (B) Drug-sensitive cells with drug introduced at t = 2 days in the absence of drug resistance. (C) Drug-resistant cells, with sensitive to resistant state transitions enabled upon addition of drug at t = 2 days. (D) Conditions same as C with epigenetic instability allowed upon drug exposure at t = 2 days. (E) Average (± standard deviations) resistant cell numbers at t = 6 days based on 2000 independent simulations. Table 1 lists parameters for each condition.

**Table 1:** Summary of the model simulation parameters. The various conditions and parameter values for each cell type, where n indicates the number of cells in a state; u is a random number parameter generating instability in the transitions; t is the time elapsed in the system; and t_max_ is the maximum time allowed in the simulations. BR: birth rate, DR: death rate and TR: transition rate.

| Condition | Time (t) | Q Cells |  |  | G-Tp, G-Tr, GS-Tp, GS-Tr |  |  | G-R, GS-R |  |  | Figure Number |
| --- | --- | --- | --- | --- | --- | --- | --- | --- | --- | --- | --- |
|  |  | BR | DR | TR | BR | DR | TR | BR | DR | TR |  |
| Control<br>No drug |  | 0 | n | 0.1*n | 2*n | n | 0.1*n | NA | NA | NA | Fig. 2A |
| Drug sensitive | t < 2 | 0 | n | 0.1*n | 2*n | n | 0.1*n | NA | NA | NA | Fig. 2B |
|  | t > 2 | 0 | 2.2*n | 0.1*n | 2*n | 2.2*n | 0.1*n | NA | NA | NA |  |
| Drug resistant<br>with stable<br>transitions | t < 2 | 0 | n | n | 2*n | n | n | NA | NA | NA | Fig. 2C |
| | t > 2 | 0 | 2.2*n | n | 2*n | 2.2*n | n | 2*n | $2.2 * n * e^{\frac{-(t-2)}{t_{max}}}$ | n | |
| Drug resistant<br>with unstable<br>transitions | t < 2 | 0 | n | n*u | 2*n | n | n*u | NA | NA | NA | Fig. 2D |
| | t > 2 | 0 | 2.2*n | n*u | 2*n | 2.2*n | n*u | 2*n | $2.2 * n * e^{\frac{-(t-2)}{t_{max}}}$ | n*u | |

Under control conditions (without drug) and minimal cell state transitions (Figure 2A), cells grow exponentially without large changes in the population of Q cells. This leads to a dynamically-balanced population where none of the cell states are dominant. In Figure 2B, drug-sensitive cell population dynamics are shown upon the addition of a drug with the condition that drug resistant populations are not allowed. Initially, there is exponential growth similar to the control conditions, followed by a decrease in all cell populations after the drug is introduced at 2 days since there are no transitions into resistant states. In control and drug sensitive cases (Figures 2A and 2B), the transition rates are minimal at all times (0.1*n). When transitions to drug resistant states are allowed the cell dynamics change accordingly (Figure 2C). Prior to addition of the drug, cells in all states grow exponentially like in Figure 2A and 2B, without any dominant cell state. Now, as drug resistance evolves, all sensitive cell state populations decrease at the expense of an increase in resistant cell state populations. This happens because sensitive cells can transition into resistant cells but the reverse cannot happen. In addition, the death rate of resistant cells decreases with time (due to acquiring resistance) but the birth rate remains the same.

Hence, unless the entire population of resistant cells die, they grow exponentially later. Due to the important role of birth and death rates of resistant cells (apart from transition rates among all cell states) in the model dynamics, we investigated their effect on population growth (Supplement section 1, Table S1). As expected, a higher birth rate of resistant cells leads to an increase in their population. Similarly, a higher death rate leads to a decrease in the resistant cell population. A high rate of transition keeps the cell population evenly distributed among the cell states (Supplementary Figure S1).

When epigenetic instability is introduced in the transition rates via a stochastic parameter (see Table 1) simultaneously with drug exposure there is a reduction in the overall population of resistant cells due to the random changes in the transition rates (Figure 2D). Since the simulations shown in Figure 2A-D represent single population trajectories, we performed a large number of simulations (2000) in order to understand the overall behavior of the model (Figure 2E). The average resistant cell population with or without epigenetic instability agree with the results from Figures 2A-D, and indicate that epigenetic instability reduces the resistant cell population size by 35%.

### Epigenetic instability in the multi-step model

The single-step basic model may not capture the complex transcriptional machinery due to PTMs and associated changes in chromatin structure, and the requirement of coordinated protein complexes to initiate gene transcription. Thus, a multi-step model was developed that considered the transition from drug-sensitive cells to drug-resistant as a multi-step lock-in process (Figures 3A and 3B). The lock-in process was implemented as a 3-step switch wherein all three events must occur to complete the transition from a sensitive to a resistant state (all switches must be On). The probability of triggering the switch is equal for all switches (*p*_1_ = *p*_2_ = *p*_3_, Figure 3B), and further, the triggering probability of each individual switch is independent of the other switches. To apply epigenetic instability, the unidirectional lock-in multi-step process is replaced by a bidirectional multi-step process (Figure 3B), where any of the three switches in an On state, can be turned Off.

**Figure 3:**
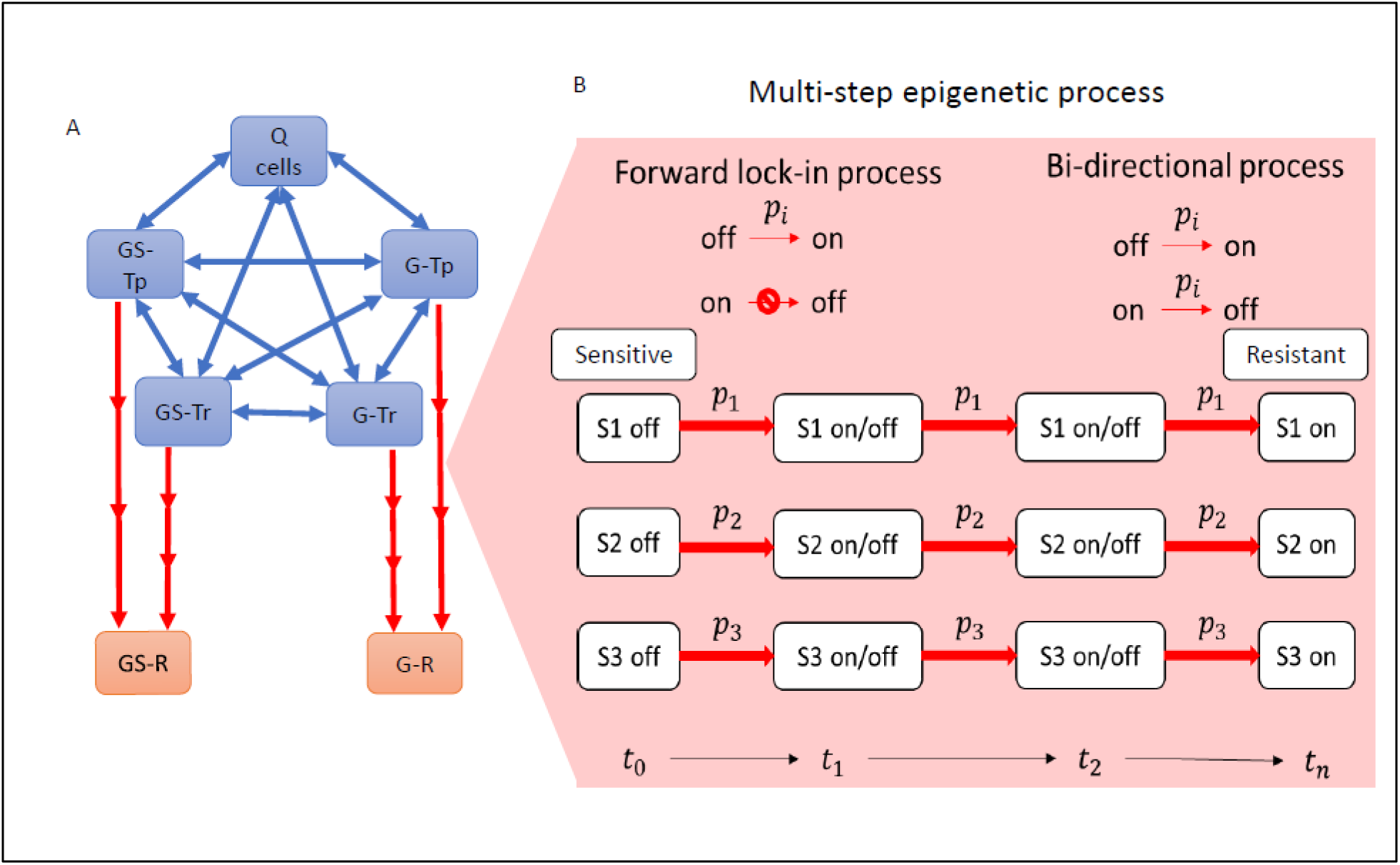
Schematic of the multi-Step model. (A) The multi-step model is similar to the basic model (see Figure 1), but transitions to drug-resistant states are multi-step processes (red arrows). Single-step processes are shown as blue-arrows. (B) The three-step transition model is shown between sensitive and resistant cell states. The forward lock-in process requires all three switches (S1, S2 and S3) to be On. The bidirectional process allows any of the three switches to go from On→Off, thus interrupting a transition to a resistant state. p_i_ indicates the probability of the event occurrence (switch turning Off→On or On→Off).

The control and drug-sensitive conditions for the multi-step model are analogous to the basic model, and thus, only the drug-resistant condition with and without epigenetic fluctuations were considered. The forward multi-step lock-in process leads to drug resistant states from both glioma stem cells and glioma cells (Figure 4A, Top panel). The parameters used to generate the simulations in Figure 4 are the same as those used in Figure 2C; however, in the multi-step model the number of resistant cells is reduced given the more complex lock-in process involved in completing a state transition. When epigenetic instability is introduced via the bi-directional process (50% probability of switch turning Off→On and vice versa), the growth of resistant cells is hampered (40% reduction) (Figure 4A, Middle panel). Moreover, high epigenetic instability (80% probability of switch turning On to Off), led to a much larger reduction (90%) in resistant cell populations (Figure 4A, Bottom Panel). As with the basic model, conducing a large number of simulations (Figure 4B) indicates that moderate epigenetic instability reduces the resistant cell population by about 60% (see Figure 4B). At high epigenetic disruption, the resistant cell population is reduced by more than 90% compared to the scenario without any epigenetic disruption.

**Figure 4:**
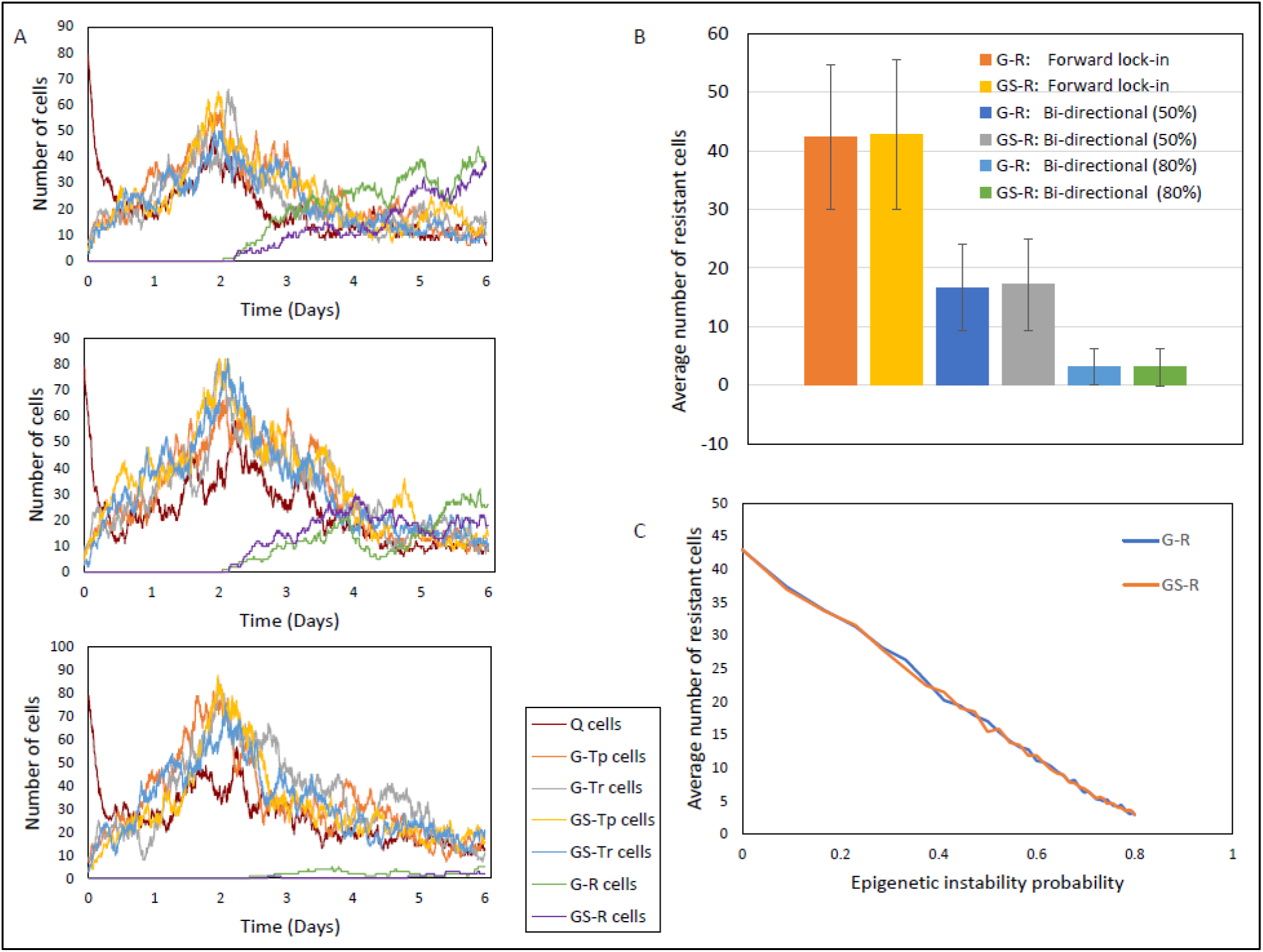
Cell population dynamics for the multi-step model. (A) Top panel: Forward lock-in multi-step process with drug introduced at t = 2 days. Middle panel: Bi-directional multi-step process with moderate epigenetic instability (50% probability of switch turning On→Off). Bottom panel: Bi-directional multi-step process with high epigenetic instability (80% probability of switch turning On→Off) (B) Average (± standard deviations) number of resistant cells at t = 6 days following 2000 independent simulations. The values 50% and 80% in the legend are the probability of switch turning On→Off. (C) The average number of resistant cells (GS-R plus G-R) as a function of increasing epigenetic instability (probability of switch turning On→Off) in a multi-step bi-directional process.

In order to determine the effect of changing levels of epigenetic instability on the resistant cell population, we performed simulations changing the probability of the switches turning On→Off. As shown in Figure 4C, there is an inverse relationship between the probability of a unit switch turning Off (a measure of epigenetic instability) and the size of the resistant cell population. At the highest tested probability of epigenetic disruption of 80%, the resistance cell numbers declined by more than 90%.

## Discussion

Anticancer drug resistance has been studied for decades with the main solution being the addition of drugs to circumvent resistance and restore cancer cell drug sensitivity [15–20]. Genetic analyses of gene mutations and expression profiles are often the basis of the drug resistance strategies that seek to identify the drug resistant-inducing genes and proteins, which may provide viable drug targets. Although the approach has appeal given the mechanistic justification, and has produced a plethora of drug combinations, these approaches ultimately fail since they do not consider intratumoral heterogeneity and innate differences in cell sensitivity to drugs, as well as cellular adaptation including the role of epigenetic reprogramming [21].

Both deterministic and stochastic mathematical models have been employed to study intratumoral heterogeneity and the emergence of drug resistance in cancer [22–27]. Among deterministic approaches, the most common use ODEs to model population growth of tumors [28]. Stochastic approaches, although computationally taxing, are generally more appropriate for modeling biological systems, which are naturally noisy and replicate the dynamics of gene transcription. In particular, discrete stochastic processes are quite relevant in modeling tumor growth as they can predict the probability of development of at least one resistant cell in a tumor [29–31]. Stochastic differential equations are used to model tumor heterogeneity as they provide a robust methodology for mechanistic modeling and the control of noise [32–33]. The stochastic modeling approach we employed is common to that used in evolutionary dynamics based on branching processes with Markovian properties that are simulated with the Gillespie algorithm [34, 35].

In this work, stochastic cell state models were developed to understand the effect of epigenetic fluctuations on tumor heterogeneity and drug resistance. Both the single-step and multi-step models show exponential growth of resistant cells upon drug exposure (Figures 2C and 4A, Top Panel). However, when epigenetic instability was introduced in either model, the resistant cell populations decreased, and were inversely proportional to the probability of the multi-step switches turning Off (Figure 4C). In the single-step model, the reduction due to epigenetic instability is about 35%. In the multi-step model, at moderate epigenetic instability (50% probability of a switch turning Off), the reduction in resistant cells was about 60% whereas at high epigenetic instability (80% probability of a switch turning Off) the reduction is more than 90%.

Both the single-step and multi-step models specify two likely epigenetic states, either transcriptionally permissive or restrictive. In the current set of simulations, there is no assigned preference to either state in terms of contributing to or mitigating drug resistant states. As such, either transcriptional state has an equal probability to transition to a resistant state. Although knowledge of specific histone PTMs that are sought for disruption may be advantageous in designing epigenetic modifier therapy, it is posed that such drug therapy cycles between transcriptionally-permissive and transcriptionally-restrictive states to maintain instability. Constant drug dosing rates are averse to instability, and lend themselves to stability in terms of the cellular programs that trigger drug resistance. Thus, although epigenetic modifiers are a route to alter histones PTM, their dosing schedules should be variable to support instability and minimize cell adaptation.

Our approach of combating anticancer drug resistance through generation of epigenetic instability may have advantages over prevailing methods. First, it reduces the complexity of ITH at least in terms of epigenetic plasticity as consisting of either transcriptionally-permissive or -restrictive cell states. This simplification – currently treated as an agnostic feature – may be tuned to repress oncogenes or to activate tumor suppressor genes by integrating the cell state model with mechanistic biochemical models. Albeit, the tuning of drug therapy to repress or activate desired genes through epigenetic modifier therapy should retain a randomness to satisfy the goal of instability. Second, a chaotic or constantly reprogramed epigenome may limit cellular adaptations including those induced by constant drug exposures. Current approaches to aggressively target one or more resistance pathways may select for alternate drug-resistant pathways to emerge. Although the resistant cells are not extinguished at 50% probability of the On/Off switch in the multi-step model, they are not proliferating or are doing so very slowly. This attribute of epigenetic instability -maintenance of this quasi-equilibrium between sensitive and drug-resistant cells - could stall or prevent implementation of alternate drug-resistant programs possibly rendering the cells susceptible to other treatments.

The proposed cell state models are theoretical and remain to be experimentally tested. Prior to that, it may be beneficial to devise a hybrid stochastic-mechanistic model to inform the most relevant experiments. For example, the task to implement epigenetic instability therapy to mitigate temozolomide resistance in brain tumor patients may be enhanced by considering synthesis of the DNA repair enzyme – methylguanine methyltransferase (MGMT) that repairs the most lethal temozolomide-induced O-6-methylguanine DNA adduct [36–38]. This would involve identifying histone PTM(s) that facilitate transcription of MGMT (a transcriptionally-permissive state) and then selecting epigenetic modifiers to prevent this process, but to do so with a sporadic dosing regimen to abate continued MGMT production whilst avoiding activation of alternate resistance mechanisms.

In conclusion, stochastic epigenetic instability models were developed that showed the potential to mitigate the emergence of drug resistant cells. The models serve as a foundation to explore detailed approaches to implement epigenetic instability therapy. It seems possible that either global or targeted epigenetic instability could be sought with combinations of epigenetic modifiers and non-constant dosing regimens. These aspirational goals will best be approached by computational models and the current cell state model is a first-step in that direction.

## Methods

### General Description of Cell State Model

The mathematical details of the cell state models and how the simulations were conducted are provided in the Supplement.

In the basic model, only single-step state transitions are allowed (see Figure 1). All state transitions are reversible except those that culminate into a drug resistant state. Once a cell is deemed drug resistant, it cannot transition to any other state. Integer cell numbers are used for each cell state to discretize cell numbers within the tumor. The system dynamics can be written in terms of a master ordinary differential equation that governs the time evolution of the probability of the system occupying a given cell state (Supplement section 2).

We incorporate stochasticity in our model by using a Monte Carlo methodology to simulate birth, death, and state transitions. As quiescent cells do not proliferate, the birth rate of Q cells is set to be zero. The death rates of quiescent cells were chosen to be a proportional to the number of cells in the state and to incorporate the effect of drug, the death rate coefficients increases after the addition of drug (See Table 1). Additionally, the Q cell population can increase or decrease due to state transitions and the transition rates were set to be proportional to the number of cells in the state from which the cell is transitioning.

For G and GS cell states, birth/death and transition rates are proportional to the number of cells in that state. Before the addition of drug, the birth rate coefficient is set higher than the death rate coefficient, which leads to exponential growth. After the addition of drugs, the death rate is set higher than the birth rate to account for the effects of drug, which leads to the decline in sensitive cells (Q cells, G and GS) populations. Moreover, the birth rate and initial death rate of the drug resistant states are kept the same as drug sensitive states. However, due to acquired drug resistance, the death rate of resistant cells decays exponentially with time.

We considered cell transitions to resistant states under two different models. In the basic model (see Figure 1), single-step one-way transitions occur from a drug sensitive state to a resistant state. In the multi-step model (see Figure 3), three uni-directional steps are required to “lock-in” a transition from a sensitive to a resistant cell state. In both the basic and multi-step models, two cases - with and without epigenetic instability – are considered.

For the simulations, we keep the initial total number of cells = 100, among which Q = 80 GS-Tp = 3 GS-Tr = 3 GS-R = 0 G-Tp = 7 G-Tr = 7 G-R = 0.

The drug is introduced in the system at time t = 2 days. In the single step model, epigenetic instability or fluctuations in the transition rate is modeled using a random number. All relevant parameters regarding each simulation is given in Table 1. To avoid complete elimination of the sensitive cell population, we apply constraints on the death rate and transition rates. If any sensitive cell state population is less than 10, then the death rate and transition rate to resistant cells is reduced to zero. All rate coefficients have units Day^−1^. A maximum time (t_max_) of 20 days is allowed during simulations. Additional details regarding the implementation of Gillespie algorithm are provided in the supplement (section 3).

## Acknowledgments

Funding from the State University of New York Empire Innovation program is acknowledged for support of Anshul Saini, Ph.D.

## Supplement

### 1. Effect of birth, death and transition rates in drug resistant cells

Different cell birth, death and transition rates were examined to assess their effect on population dynamics (Table S1). A lower birth rate (1.5*n) led to a smaller resistant cell population in comparison to a higher birth rate (2.5*n) (Figure S1A-B). To illustrate the effect of drug-induced time dependence on the death rate of resistant cells, two different death rates (e^−(t− 2)/10^ and e^−(t− 2)/100^) were used. In the first case (e^−(t− 2)/10^), the death rate decreased rapidly leading to an exponential rise of resistant cell population, whereas in the second case (e^−(t− 2)/100^), the death rate decreased slowly, and thus resistant cells died at a faster rate (Figure S1C-D).

We next considered cell states under two conditions; either having no transitions or having high transition rates (2*n). In the absence of transitions, there is no resistant cell population as resistant cells arise from sensitive cells. Additionally, the cell state trajectories remain independent of each other resulting in large disparity in the number of cells occupying different cell states (Figure S1E). In the case of high transition rates, resistant cells dominated the cell population. However, allowing transitions between cells states changed the behavior of the system by allowing the cells to move from more populated states to lesser-populated states (Figure S1F). Hence, most of the cell state trajectories remained close to each other.

**Table S1:** Summary of model simulation conditions with alternate birth, death, and transition rates. n indicates the number of cells in a state; t is the time elapsed in the system. BR: Birth rate, DR: Death rate and TR: transition rate.

| Condition | Time<br>(t) | Q Cells |  |  | G-Tp, G-Tr<br>GS-Tp, GS-Tr |  |  | G-R, GS-R |  |  | Figure<br>Number |
| --- | --- | --- | --- | --- | --- | --- | --- | --- | --- | --- | --- |
|  |  | BR | DR | TR | BR | DR | TR | BR | DR | TR |  |
| Low BR<br>of G-R,<br>GS-R | $t < 2$ | 0 | n | n | $2*n$ | n | n | NA | NA | NA | Fig.S1A |
| | $t > 2$ | 0 | $2.2*n$ | n | $2*n$ | $2.2*n$ | n | $1.5*n$ | $2.2*n * e^{-(t-2)/20}$ | n | |
| High BR<br>of G-R,<br>GS-R | $t < 2$ | 0 | n | n | $2*n$ | n | n | NA | NA | NA | Fig.S1B |
| | $t > 2$ | 0 | $2.2*n$ | n | $2*n$ | $2.2*n$ | n | $2.5*n$ | $2.2*n * e^{-(t-2)/20}$ | n | |
| Low DR<br>of G-R,<br>GS-R | $t < 2$ | 0 | n | n | $2*n$ | n | n | NA | NA | NA | Fig.S1C |
| | $t > 2$ | 0 | $2.2*n$ | n | $2*n$ | $2.2*n$ | n | $2*n$ | $2.2*n * e^{-(t-2)/10}$ | n | |
| High DR<br>of G-R,<br>GS-R | $t < 2$ | 0 | n | n | $2*n$ | n | n | NA | NA | NA | Fig.S1D |
| | $t > 2$ | 0 | $2.2*n$ | n | $2*n$ | $2.2*n$ | n | $2*n$ | $2.2*n * e^{-(t-2)/100}$ | n | |
| No<br>Transition<br>rate | $t < 2$ | 0 | n | 0 | $2*n$ | n | 0 | NA | NA | NA | Fig.S1E |
| | $t > 2$ | 0 | $2.2*n$ | 0 | $2*n$ | $2.2*n$ | 0 | $2*n$ | $2.2*n * e^{-(t-2)/20}$ | 0 | |
| High<br>Transi-<br>-tion rate | $t < 2$ | 0 | n | $2*n$ | $2*n$ | n | $2*n$ | NA | NA | NA | Fig.S1F |
| | $t > 2$ | 0 | $2.2*n$ | $2*n$ | $2*n$ | $2.2*n$ | $2*n$ | $2*n$ | $2.2*n * e^{-(t-2)/20}$ | $2*n$ | |

**Figure S1:**
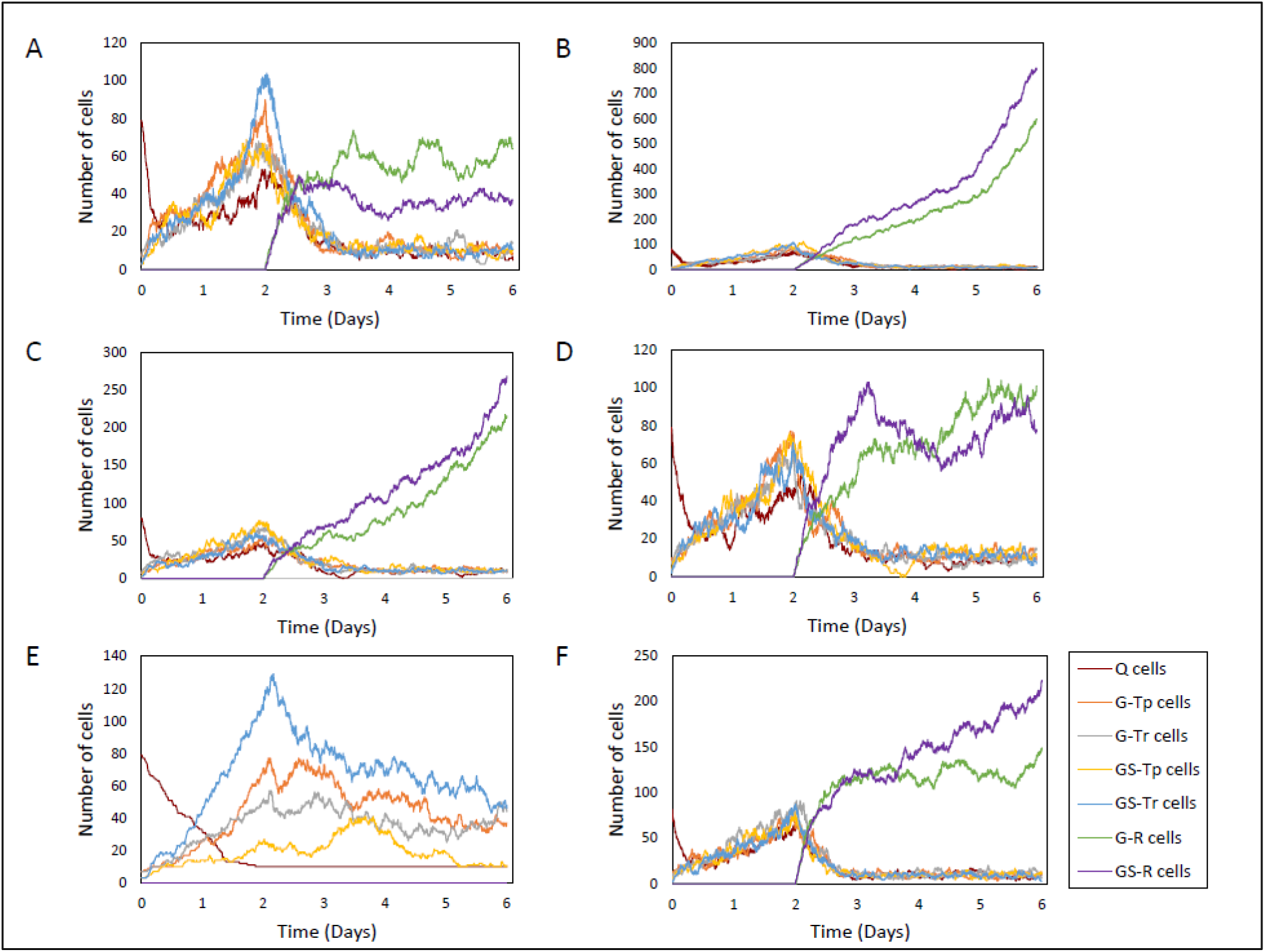
Effects of changing birth/death/transition rates on cell population dynamics. (A) Simulation for low birth rate of resistant cells (1.5*n). (B) Simulation for high birth rate of resistant cells (2.5*n). (C) Simulation for low death rate of resistant cells (2.2*n* e^−(t− 2)/10^). (D) Simulation for high death rate of resistant cells (2.2*n* e^−(t− 2)/100^). (E) Simulation without any transition. (F) Simulation with high transition rate (2*n).

### 2. Mathematical description of the model

As stated before, in our model, there are seven cell states, which are denoted as a row vector {*n*_1_, *n*_2_, ‥ , *n*_7_} (*n*_*i*_ represent the number of cells in the *i*^th^ state). We assume continuous time Markovian dynamics for cell proliferation, which means that the future evolution of the system can be predicted from the current state alone (memoryless). The process of birth, death, and transition between any cell states *i* and *j* can be written as

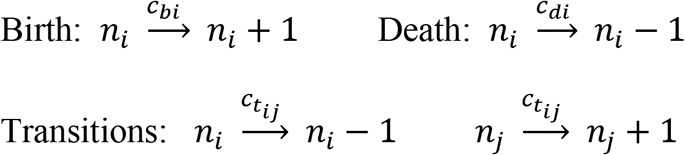

where *c*_*bi*_ and *c*_*di*_ is the birth and death rate coefficient in *i*^*th*^ cell state, respectively. 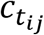 is the transition rate coefficient regarding the transition from *i*^*th*^ → *j*^*th*^ cell state.

All of the above processes can be written in terms of transitional probabilities (Figure S2) as

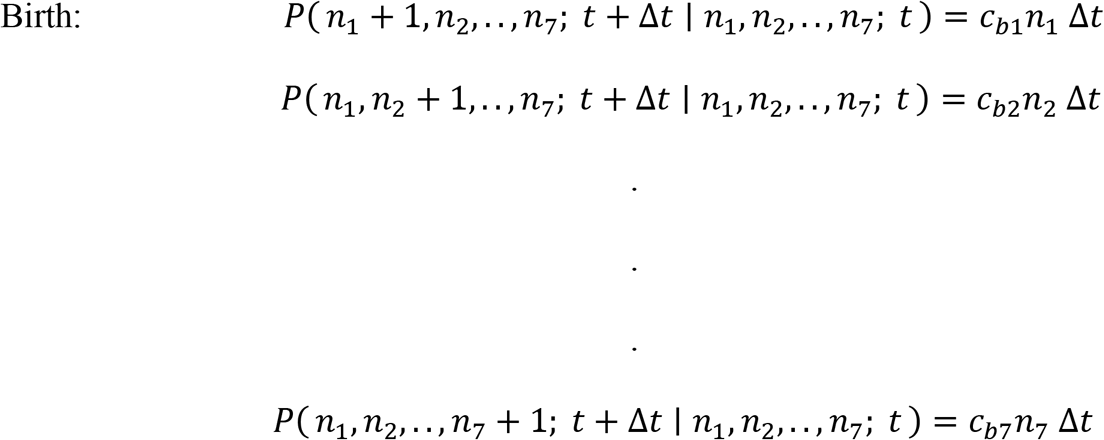

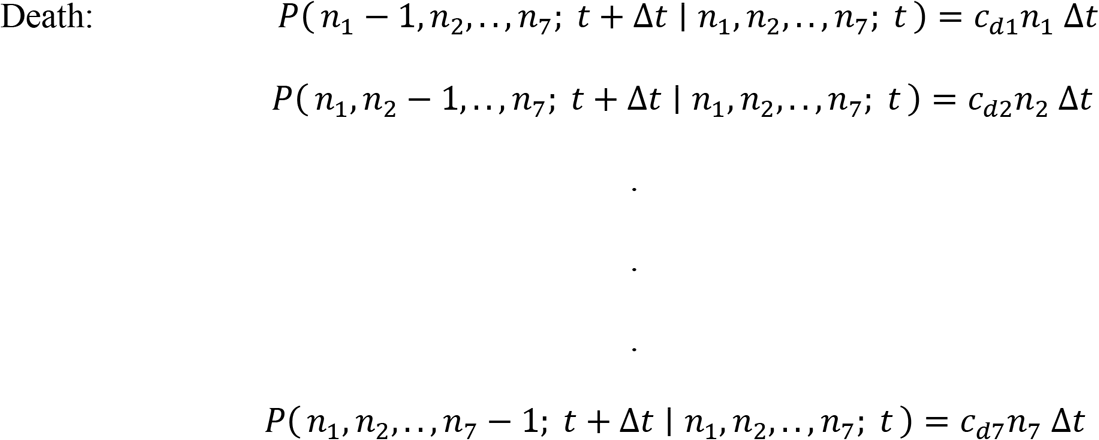

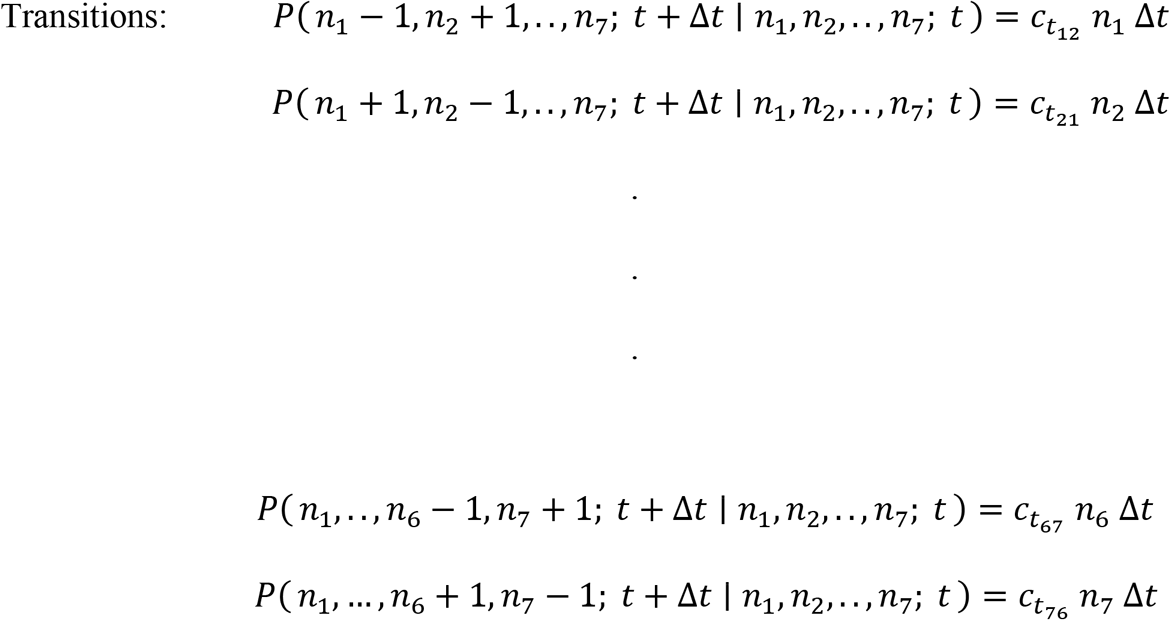

In stochastic systems, time evolution is given by a master equation, which is derived by supposing that the system is in state {*n*_1_, … , *n*_6_, *n*_7_} at time t, and based on the transitions defined above, we can write the probability distribution of the state {*n*_1_, … , *n*_6_, *n*_7_} at time *t* + Δ*t* as

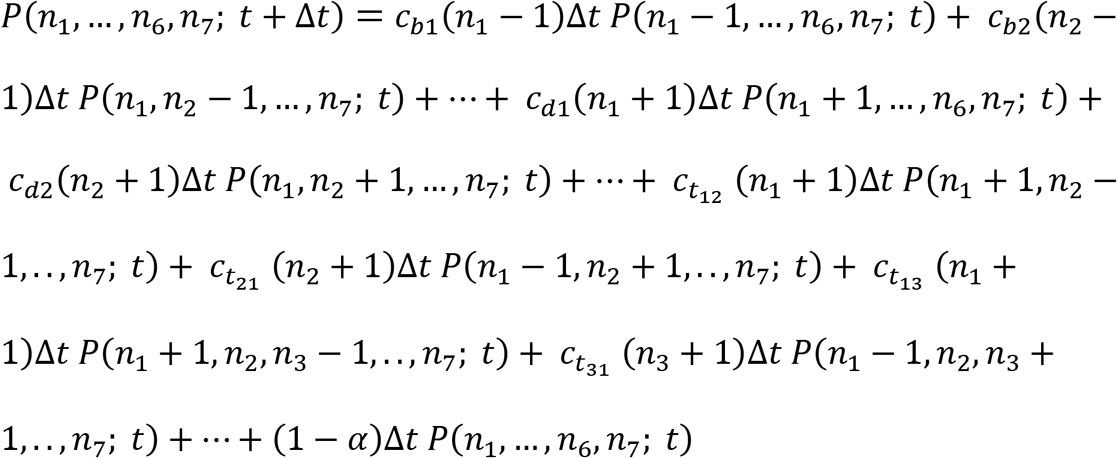

where *α* is the sum of all transition probabilities from other states to the state {*n*_1_, … , *n*_6_, *n*_7_}. The above equation can be written in a succinct form as

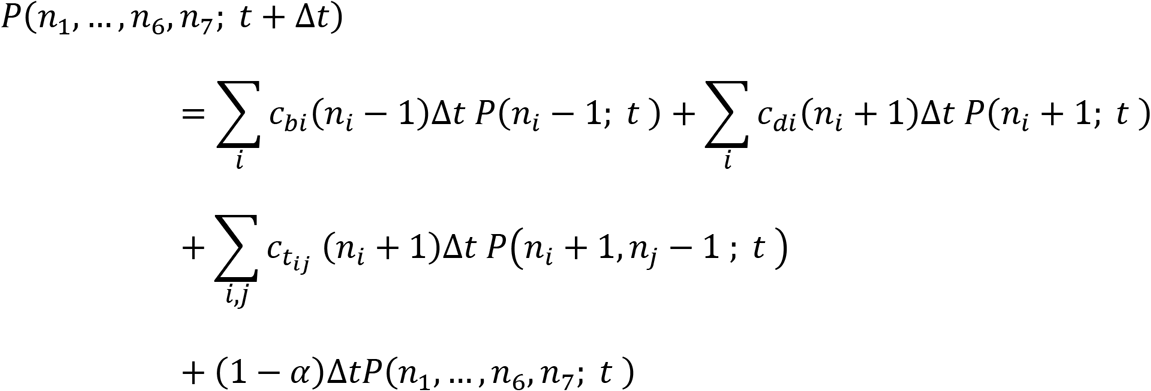

Applying a few algebraic manipulations yields the master equation as

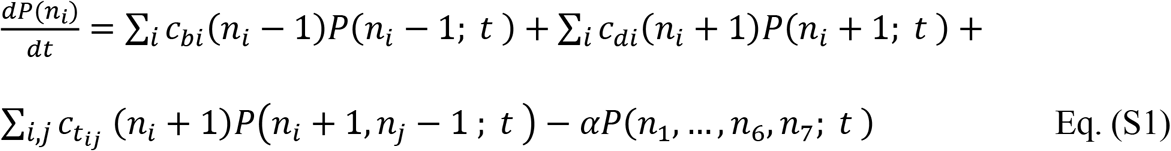

The first term corresponds to the birth, the second term to death, the third term represents outward and inward transitions from and to a particular state, and the last term is the probability of none of the above events occurring. Both birth and death rates are proportional to the number of cells in the corresponding state. The transition rate is proportional to the number of cells in the initial state (*n*_*i*_). The coefficients are set according to the birth rates, death rates, and transition rates between the states.

Once the initial values of (*n*_*i*_) and coefficients (*c*_*bi*_, *c*_*di*_, 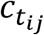) are known, the stochastic time evolution of the system can be evaluated using the master equation.

**Figure S2:**
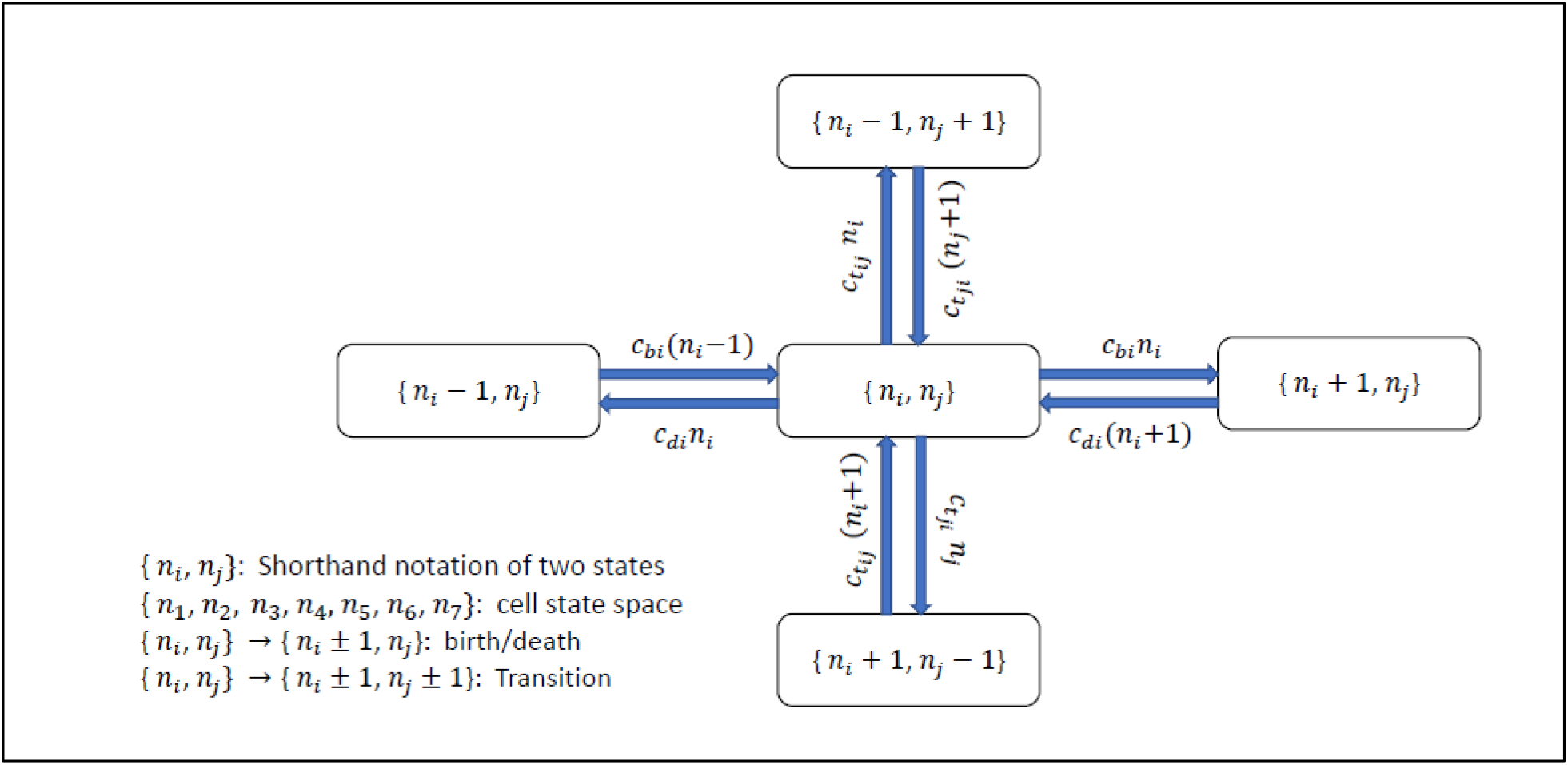
Illustration of the flow of cell state probability within the cell state space through the process of birth, death, and transition. *n*_*i*_ Indicates the number of cells in the *i*^*th*^ state. *c*_*bi*_, *c*_*di*_ and 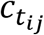 are birth, death and transition rate coefficient, respectively.

### 3. Simulations

To simulate the system, the Gillespie algorithm was used, where individual trajectories are simulated instead of the whole probability distribution. The algorithm assumes the processes to be Markovian and a time step (*τ*) is used based on the probability of the event occurrence (Poisson distribution). To find *τ*, we define the propensity function as

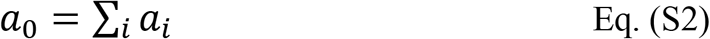

where the probability that any given event will occur per unit time is *a*_*i*_. The time before the next reaction occurs is a random variable with distribution

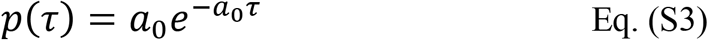

We can generate an exponentially distributed τ (time to next reaction) by

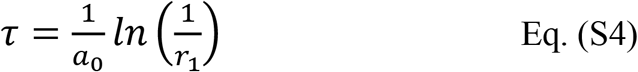

where, *r*_1_ is a uniformly distributed random number in (0, 1). We used another random number to select which of the reactions (based on their weight) is going to take place. If event 1 is selected: *n*_1_ → *n*_1_ + 1. If event 2 is selected: *n*_1_ → *n*_1_ − 1 and so on. We update the number of cells in each cell state and repeat the process until the desired time or maximum cell population is reached. The code is available on GitHub (https://github.com/anshulsa/Cell_state_transition.git).

